# DNA Circle-sequencing lowers the single molecule sequencing error threshold and identifies ultrasonication as a source of DNA damage

**DOI:** 10.1101/2025.07.30.667586

**Authors:** Stephan Baehr, Jean-Francois Gout, Lauren Reyes, Haimanti Ray, Michael Lynch

## Abstract

DNA mutation is the ultimate source of all heritable genetic and phenotypic variation. On human timescales, DNA mutation results in the evolution of antibiotic resistance, viral resistance to vaccines, the emergence human cancers, and more. Precise measurement of DNA mutation is therefore desirable to rapidly detect and analyze low frequency mutations in a population of cells. Precise measurement of DNA mutation by high throughput sequencing has been hindered by sequencing error rates, which occur at rates 1×10^−2^ to 1×10^−3^ per base sequenced. Previous work on circle sequencing had pushed a putative sequencing error rate down to roughly 2.8×10^−4^ per base for genomic yeast DNA in the absence of DNA repair enzymes, where the expected number is 4.7×10^−9^ per base per generation. Through revision of the assay, we are now capable of taking 125ng of genomic DNA and obtaining mutation rate estimates with a sequencing resolution floor of 2×10^−7^ per base in the absence of repair enzymes, a resolution improvement of over 3 orders of magnitude. In practice, circle-seq recovers some of the mutation spectrum of mismatch-repair deficient *E. coli*, although some signature of C → T and G → A errors remain present. Curiously, it calls a mutation rate of 2×10^−9^ per base per generation, lower than expected, underscoring the difficulty of directly comparing mutation rates per base to MA mutation rates per site per generation. This protocol readily recognizes Covaris ultrasonication as mutagenic in library preps, with a mutation spectrum dominated by G:C → A:T, G:C → C:G, and G:C → T:A errors. The protocol is either detecting *in vivo* DNA damage that has yet to be repaired, is significantly biased in its ability to detect mutations, or is not yet sensitive enough to detect mutation rates of MMR-*E. coli* cells. The protocol may have some use in measuring mutation abundance from tissue samples, and *in vitro* sources of DNA damage. The protocol also highlights some biases inherent to single-stranded DNA (ssDNA) cyclization, including a 10-bp periodicity which echoes the constraints of dsDNA cyclization.

## 1 Introduction

DNA mutation is the basis of all heritable biological variation. Some genetic variation gives rise to beneficial traits, most mutations are neutral or nearly neutral[6, 13], and some mutations result in severely deleterious phenotypes[1]. DNA mutation is known to affect evolutionary trajectories of phenotypes over millions of years, but advances in the last century have shown that evolution can occur over hours, days, or years[27, 31, 15, 16, 42]. For example, the emergence of organisms resistant to herbicides, pesticides, and antibiotics threaten great triumphs of biotechnology and medicine[4, 33]. Rapid evolution of the common cold and novel viruses such as COVID-19 decrease human productivity[43] and have proven their ability to constrict our global economy. Accumulated mutations in our somatic cells cause cancer[14], help cancer to evade anticancer treatments[5], and broadly degrade the efficiency of bodily functions in aging[41]. Pinpointing novel mutations in populations of cells is therefore of particular interest to the study of diverse human-related subjects, from viral evolution to personalized medicine and predictive medicine.

Precise measurement of DNA mutations using high-throughput sequencing is hindered by the error-prone nature of sequencing machines, which generally have error rates between 1 in 100 bases or 1 in 1000 bases mis-called. It is difficult to measure things that occur once in a million bases, when background noise signal occurs every thousand bases[34]. To get around this limitation, researchers have employed several techniques. Before DNA was even identified as the genetic material on which mutations arise, the fluctuation test[27, 24**?**] was accurately estimating mutation rates, and remains a functional technique[22, 42]. Other reporter assays have been used since the earlier years and are numerous[38]. The mutation accumulation experiment[42, 28] can measure any mutation rate fixes mutations at 100% frequency within lines. Variations of the mutation accumulation experiment have been used to estimate somatic cell mutation rates[32], along with trio studies of germline mutations in organisms with large genomes[32, 3]. The use of laser-capture microdissection additionally provide somatic mutation estimates by duplicate or triplicate sequencing of nearby samples[18, 7]. SMRT-sequencing has also been used to measure somatic mutation and DNA damage within tissues with success[30], particularly by sequencing both sense and antisense strands of DNA. DNA barcoding[39] paired with ultra-deep sequencing is employed to detect rare mutations, and Duplex-seq[35] is yet another method developed to get around high sequencing error rates. Though all of these methods work, all have their drawbacks, including high labor costs, low throughput, low genome coverage, or some combination thereof.

One particular method developed in 2013[26], “Circle-sequencing”, appeared to have exemplary attributes with regard to labor cost and throughput. When it was published it claimed the lowest potential resolution floor for sequencing, by employing tandem repeats and DNA repair enzymes. These innovations sought to ameliorate DNA damage induced during the DNA extraction and library preparation steps prior to sequencing. However, we found that the error rate observed by this publication, 7.6 × 10^−6^, was replicated for organisms known to have mutation rates several orders of magnitude lower than this reported rate, for example mismatch-repair deficient (MMR-) or wild-type *E. coli* strains.

The allure of the technique remains, if somehow the problem of *in vitro* DNA damage could be resolved. In effect, a person could take a tumor biopsy, run a standard DNA extraction, and then a library prep, all of which in principle could be, and are, highly automatable. This technique, if proven, would provide a relatively economic and fairly rapid (2-week turnaround is very feasible) means of calling all the extant mutations of a tumor, and if a comparison tissue is given, a mutation rate per distance or per unit time may be estimated. To demonstrate the efficacy of the DNA Circle-seq technique, we have chosen to reproduce known mutation rates and known, distinctive, and unmistakable mutation spectra[25, 42, 2] in *E. coli*. It has proven challenging to equate mutation rate per base measurements to mutation rates obtained by MA experiments, the gold standard of mutation estimation technology; the mutation spectrum however, should translate well.

We describe a technique of measuring mutations and mutation rates, which significantly refines and revamps the original circle-sequencing protocol first employed in 2013 and expounded upon in RNA applications in 2017 and beyond [17, 8, 9]. DNA RCA demonstrates some ability to detect the mutation spectrum of MMR-*E. coli*, though it also elicits an additional spectrum of G and C damage, a signature of oxidative damage known to occur *in vitro* and perhaps *in vivo*. Circle sequencing’s mutation rate estimate of MMR-is on the order of 2 × 10^−^7 per base, or 3.68 × 10^−9^ mutations per base per generation, which substantially differs from MA experiments, for which we cannot account. We demonstrate that Covaris ultrasonication induces DNA damage, and increases mutation rates to roughly 2 × 10^−5^ mutations per base by the library preparation protocol used, on the order of 100-fold. The assay has been unable to fully recapitulate the mutation spectrum of MMR-*E. coli*, for either *in vivo* or *in vitro* complicating factors. We also note that this assay is suitable for the study of the biases inherent to ssDNA cyclization, and dsDNA cyclization, with lower throughput. We describe the distinctive 10-bp periodicity of cyclization efficiency, and note the potential of GC content bias in the efficiency of loop formation.

## 2 Methods

### 2.1 Strain information

The *E. coli* used in the study is a descendant of the Foster lab’s 2012[25] mutation accumulation experiment. The Wild type “WT” genomic background is *E. coli* K-12 str. MG1655, and “MMR-” or mismatch-repair deficient, is derived from the WT strain by a mutL deletion, which causes *E. coli* to lose the ability to perform DNA replication mismatch surveillance. This results in a 100 to 150-fold increase in transition mutations A:T → G:C, and G:C →A:T. Strain 406-1, also known as MMR-L10-A1, “S-”, an *E. coli* clone that emerged as a result of a long-term evolution experiment on the MMR-genetic background for 1000 days in the Lynch lab[42]. A mutation accumulation experiment has been run on all strains, identifying their significantly different mutation rates. 406-1 had the highest mutation rate measured at roughly 4 × 10^−7^bps/site/gen, or roughly 2000-fold of the WT *E. coli* mutation rate.

### 2.2 Growth conditions and DNA extraction

*E. coli* were first grown overnight in liquid culture, 37°C, 200rpm, (1x LB Broth, Miller formulation) in borosilicate glass 10mL tubes with loosely fitting metal caps from frozen stocks. The overnight culture was then serially diluted (100uL culture to 900uL 1x PBS) in a biosafety cabinet to a range of dilutions around 1 ×10^−7^ and 1 ×10^−8^. Either 20uL or 50uL from each dilution was used to seed 10 10mL tubes, with the aim of reaching a dilution rate of 0.5 bacteria per inoculum. When the correct dilution is hit, about half of the 10 tubes inoculated will become turbid overnight from *E. coli* growth, and half will remain clear. Any ratio of 0.5 of 10 tubes or lower with ( E. coli) growth is indicative of growth from a single cell, under the assumption of a Poisson distribution of a discrete variable. *E. coli* were grown by two methods; 1. overnight for 10 hours, from 10pm to 10am, at 37°C, shaken at 200rpm; 2. for 24 hours, resulting in some degree of stationary phase post-growth. Three ideal tubes from the serial dilution sets are chosen as biological replicates. Colony Forming Units (CFU’s) are taken from the 10mL tubes, and the rest of the culture is used for DNA extraction (Wizard Genomic DNA Extraction Kit, Promega), with the following protocol modifications: cells were heated only to 65°C for 5 minutes with intermittent vortexing every minute to lyse cells, instead of 80°C recommended by the protocol. Also, due to somewhat overloading the extraction protocol, 400uL instead of 200uL of the protein precipitation salt was used, and a 10min spin of 16,000 x G @ room temperature was inserted post-precipitation to form a more solid pellet. The protocol otherwise followed the manufacturer specifications.

Genomic DNA (gDNA) was run through a gDNA cleanup column (Zymo Research) and eluted in autoclaved milli-q h20, pH 8, and stored at least overnight in a −80°C freezer. DNA concentration was then measured via Qubit 4.0 with the 1x dsDNA HS kit; occasionally 1:10 dilutions were required to obtain a DNA concentration measurement, since the kit is most accurate at concentrations 10-40ng/uL. Importantly, we found the 2-minute incubation before loading into the Qubit machine, as per the manufacturer’s instructions, to be critical for measuring DNA concentration accurately and reproducibly over time and across samples. 125ng of gDNA was then used as the input of the mu-seq protocol, see Figure 1 and supplemental methods for a full protocol. Briefly, the DNA is first fragmented with MNase (NEB), an enzyme with both endonuclease and exonuclease activity. Importantly, MNase is sensitive to degradation by vortexing; for a proper dilution, vortexing must be avoided. Instead, a 2uL to 60mL dilution was obtained by adding the MNase to a bottle of water, pH 8, and inverting 20 times to mix before being put on ice. The enzyme does degrade in water over time, but it is not relevant over the time period of an afternoon when on ice.

**Figure 1.**
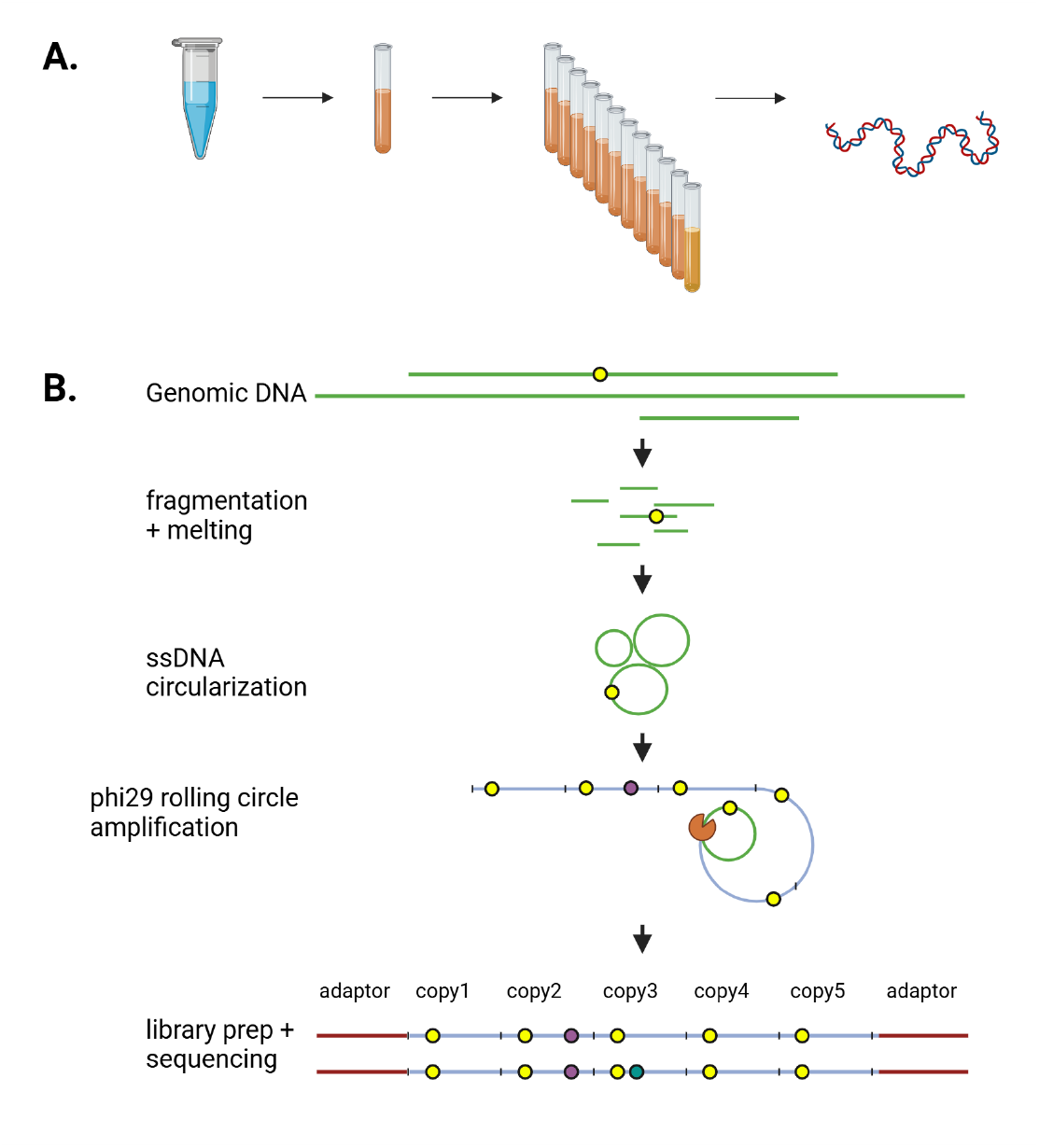
Circle-Seq Protocol. A. Frozen *E. coli* stocks are thawed and grown 24 hours. The stationary phase culture is serial diluted. After 12 or 24 hours growth in a 10mL tube, DNA is extracted. B. The DNA circle-seq protocol begins with genomic DNA, which is fragmented. After fragmentation, the DNA is end-repaired, melted, circularized, and then undergoes rolling circle amplification. True mutations (yellow circles) are present at the beginning. These mutations end up on some circles, which when amplified, produce a mutation in the exact same location in tandem repeats. Polymerase errors (purple circles) emerge, as do sequencing errors (blue circle). However, in contrast to true mutations, these errors occur randomly, and can be bioinformatically discerned from one another. The only risk to the protocol are *in vitro* DNA damage events that occur prior to rolling circle amplification, as they will have the same repeat pattern.

After fragmentation and size verification, the DNA is treated with polynucleotide kinase (PNK) due to the shearing mechanism of MNase leaving 3’ phosphates. The DNA is then melted with pH rather than heat; pH 12.5 is sufficient to turn double-stranded DNA into single-stranded DNA, which, if it is mutagenic, is less so than high temperature (95°C). After DNA melting, the DNA is circularized to itself via RNA ligase I. No matter the concentration tested in our efforts, self-ligation appears far more likely to occur than oligomer-ligation, although we must concede it is likely that some degree of this is occurring, which we fail to detect. Circularization is followed by an exonuclease digest, which destroys non-circularized DNA. After cleanup of the circularized DNA, Rolling circle amplification is completed with phi29 and random hexamers.

Upon completion of the rolling-circle amplification portion of the circle-seq protocol, the existence of visible fine long strands of DNA in the PCR tube are a strong indicator of a successful prep. After cleaning the DNA with a Zymo Oligo Clean and Concentrator, the DNA was assessed for quality control: fragment size, concentration, 260/230 ratio on the nanodrop. Routinely, 1-5 micrograms of DNA are produced from the protocol initiation of 125ng, depending on the input DNA template and the fragment size; smaller fragment sizes produce less. The DNA was either shipped a sequencing core for library prep, or alternatively library construction was completed in-house. If in-house, we used the covaris ultrasonicator to shear the DNA to proper library sizes, followed by standard library prep following the NEBnext Ultra II protocol, save that we covaris sonicate our DNA to acquire the proper insert sizes for sequencing. Sequencing has been performed on the Illumina Hiseq 2500, Illumina Novaseq 6000, and BGI Americas sequencing platform DNB-seq. BGI Americas uses a pcr-free library prep kit for input amounts >1 microgram, which all libraries have met.

### 2.3 Changes to Circle-sequencing

To begin, we looked at the DNA extraction and chose to avoid high temperature and phenol-chloroform extraction[10]. We also removed all high temperature steps utilized to inactivate enzymes. The input DNA amount was reduced from 2.5 micrograms to 250ng, and then later to 125ng. The DNA fragmentation was changed from sonication to enzymatic fragmentation with MNase. We have improved the fragmentation reaction to increase reliability/reproducibility of the protocol, by changing the dilution step of the manufacturer concentration from 2uL/60mL, reducing pipettor error. The original protocol[26] calls for a gel size selection after fragmentation; we were able to skip this. The DNA was melted by pH instead of by temperature. The circularization step enzyme was changed from CircLigase II to RNA ligase I, although 3 microliters of ligase are added instead of 1 microliter; the circularization temperature dropped to 25°C from 60°C. From the circularization reaction we removed the DNA repair enzymes FPG and UDG. The rolling circle amplification (RCA) temperature “primer annealing” step @ 65°C was also removed, being found to be irrelevant. The RCA phi29 polymerase concentration was doubled, and the DNTP concentration increased 2.5x. Also, we noticed in the 2x annealing buffer the troubling addition of too much EDTA; This was reduced from 1mM to 1 uM, significantly increasing yield. Post RCA, we originally sequenced the reads with the Hiseq 2500 with paired-end 250bp reads, and additionally to the Novaseq 6000. However, in an effort to increase the flexibility of the assay to more platforms with higher yields, we modified the assay for PE150, and used Beijing Genomics Institute’s DNB-seq T7 platform.

The end result is a leaner protocol that takes 1/20th the input DNA, takes half the time to execute, produces micrograms of DNA for sequencing, and most importantly, significantly reduces the sequencing error rate encountered by the protocol.

### 2.4 Cell division count

Colony Forming Units (CFU’s) were determined from 12 hour or 24-hour cultures by standard techniques, diluting to the 10^−6^ and 10^−7^ plates. Colonies were counted and an average was obtained. On average, 28 cell divisions were responsible for the growth of all samples except the DNB-seq samples, which had 33 cell divisions of growth (Supplemental Information). However, because of filtering protocols of the Data Analysis section, the first 5 cell divisions are discarded by our protocol. Thus, we divided our base-pair substitution per base sequenced (bps/base) rates by 23 and 28, respectively, in order to obtain an estimate of bps/base/generation, which we deploy as a potential estimate of bps/site/generation.

### 2.5 Data Analysis

We modified a pipeline previously used to measure RNA-seq reads generated by circle sequencing. Briefly, this pipeline started by identifying repeats within each read based on sequence similarity (minimum repeat size, 30 nucleotide (nt); minimum identity between repeats, 90%). Then, a consensus sequence of the repeat unit was built by summing the quality score of all four possible base calls (A, T, C, or G) from the repeats at each position and retaining the one with the highest total quality score. The next step consisted of identifying the position in the consensus sequence that corresponded to the 5’ end of the DNA fragment (because phi29 dna polymerase is randomly primed, the concatemer of DNA—and therefore, the read sequence—can start anywhere on the circularized ssDNA). This was carried out by searching for the longest continuous mapping region in a BLAST mapping of a tandem copy of the consensus sequence against the reference genome. The consensus sequence was then reorganized to start from the identified ligation point (that is, the 5’ end of the original ssDNA fragment). This reorganized consensus sequence was then mapped against the genome with TopHat (version 2.1.0 with bowtie 2.1.0), and all nonperfect hits went through an algorithm of refining the search for the location of the ligation point before being mapped again. Finally, every mapped nucleotide was inspected and must pass a number of thresholds to be retained: (i) The mapped nucleotide must be supported by at least three repeats from the original sequence reads, (ii) all repeats must support the same base call, (iii) the sum of base call qualities at this position is above 100, (iv) the nucleotide must be more than 5 nt away from the end of the consensus sequence (to minimize false-positives induced by mapping errors), and (v) the nucleotide must also be at a genomic position covered by at least 20 reads and with less than 5% of these reads supporting a base call different than that of the reference genome (this allows for filtering out polymorphic sites). For each read containing at least one mismatch passing these thresholds, sequences corresponding to all possible versions of the position of the ligation point were generated and mapped against the genome with TopHat. If at least one of these sequences finds a perfect match, then the original read is discarded. This last test only removes a small fraction of the error-containing reads (typically less than 5%), but it ensures that errors in calling the position of the ligation point cannot produce false-positives.

Every mapped nucleotide that passes all these thresholds was considered as a potential event of DNA mutation. Since all experiments were run in triplicate, two final tests of independence of mutations were run. First, we noticed that there was a particular excess of mutations arising in the first two and last two nucleotides of the repeat; this is likely due to mis-mapping, or less likely DNA damage; regardless, these mutation calls were omitted. Secondly, if a mutation is found in the exact same place in more than one replicate, it is discarded as a mismapping error. Iterating upon that theme, several regions of the *E coli* genome were blocked out in full, due to their highly repetitive nature which lead to numerous false positives shared between multiple samples. For example, the ribosomal RNA locus has tandem repeats of the same sequence with some minor variation; when the repeat length is between 20 and 50 bases, it is inevitable that some mismapping will occur along the locus and distort an estimate of mutation rate. The full list of masked regions are as follows: masked the genome for the following regions: 19860 to 20120, a transposase; 223901 to 228767, rRNA region; 270331 to 275096 transposase and phage regions; 729708 to 737736, highly repetitive region. The total DNA mutation rate was calculated as the number of unique mismatches divided by the total number of mapped nucleotides that passed all quality thresholds. Because some mutations will arise “early” in the growth of the sample, and are known as jackpot mutations, we chose to assume via the infinite sites assumption that any mutation identified more than once within a single sample arose from a single mutation event.

### 2.6 Mutation Annotations

Mutation annotations were generated by conventional means, using SNPEFF with a custom built *E. coli* database to NCBI Nucleotide genome assembly U00096.2. Selection was detected by the standard equation 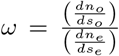, where *ω* is the dN/dS ratio[20], or the ratio of ratios of the fraction of nons-ynonymous to synonymous mutation calls observed, divided by the random non-synonymous to synonymous mutation ratio, the null expectation. The expected ratio of non-synonymous to synonymous sites was estimated from simulation of the mutation spectrum.

### 2.7 Mutation simulations

Mutation simulations were run with via an in-house python script which generates random mutations relative to a given mutation spectrum. The mutation spectrum of the liquid MA is relatively close to neutral evolution[23, 28, 2] and therefore was used as a best means of generating random mutations, rather than simply counting the number of non-synonymous and synonymous sites in the *E. coli* genome.

### 2.7 Fragment size periodicity detection

A basic histogram is sufficient to parse the relationship between fragment size and count. When multiple samples vary in sequence depth, the average count between lines creates a useable graph with potentially high error bars. A modified strategy was used to normalize the relative sequence depth between samples. fragment length counts were normalized for each sample to the total number of reads in each sample. These fractions ranged from 1 ×10^−^7 to 1 ×10^−^1. Log-transformed bar graphs in R did not behave well, so all values were multiplied by 1 ×10^7^ to create positive integers which reflect the relative abundance of each fragment length.

## 3 Results

The method illustrated in Figure 1 was used on *E. coli* strains, and is expounded upon in further detail in the Methods section. To begin, echoing the original 2013 publication, we sought demonstrate how many repeats may or ought to be used to detect mutations. From paired-end 250bp reads, the maximum total sequence length of a read is 500bp. This is important to keep in mind with regard to the number of tandem repeats needed to detect a mutation; as repeat counts increase, only smaller repeats will possibly provide 4 full repeats for example. There is a trade-off in the size of the starting fragments chosen for the assay, and the eventual yield of “unique” bases. We chose the final set of samples processed, MMR-*E. coli* procured via DNB-seq, to investigate this question. Figure 2a calls mutations from reads corresponding to 2, 3, or 4 repeats from the triplicate samples, and counts the mutations detected relative to the number of unique bases called. Figure 2a demonstrates that the mutation spectra and summed mutation rates vary per base, depending on the number of repeats used to call mutations. The DNB-seq samples using 3 repeats calls the lowest mutation rate. When compared to the mutation spectrum of Figure 2b, relative to the known mutation spectrum of *E. coli* grown in near-identical liquid culture conditions, it is clear that none of the DNB-seq repeat counts fully recapitulate the purple Liquid MA spectrum. However, as to which one is closes, the 3-repeat analysis trikes closest to the mark. In all cases transitions are more common than transversions, but the relative ratio of AT>GC and GC>AT mutations is modestly closer to the baseline expectation.

**Figure 2.**
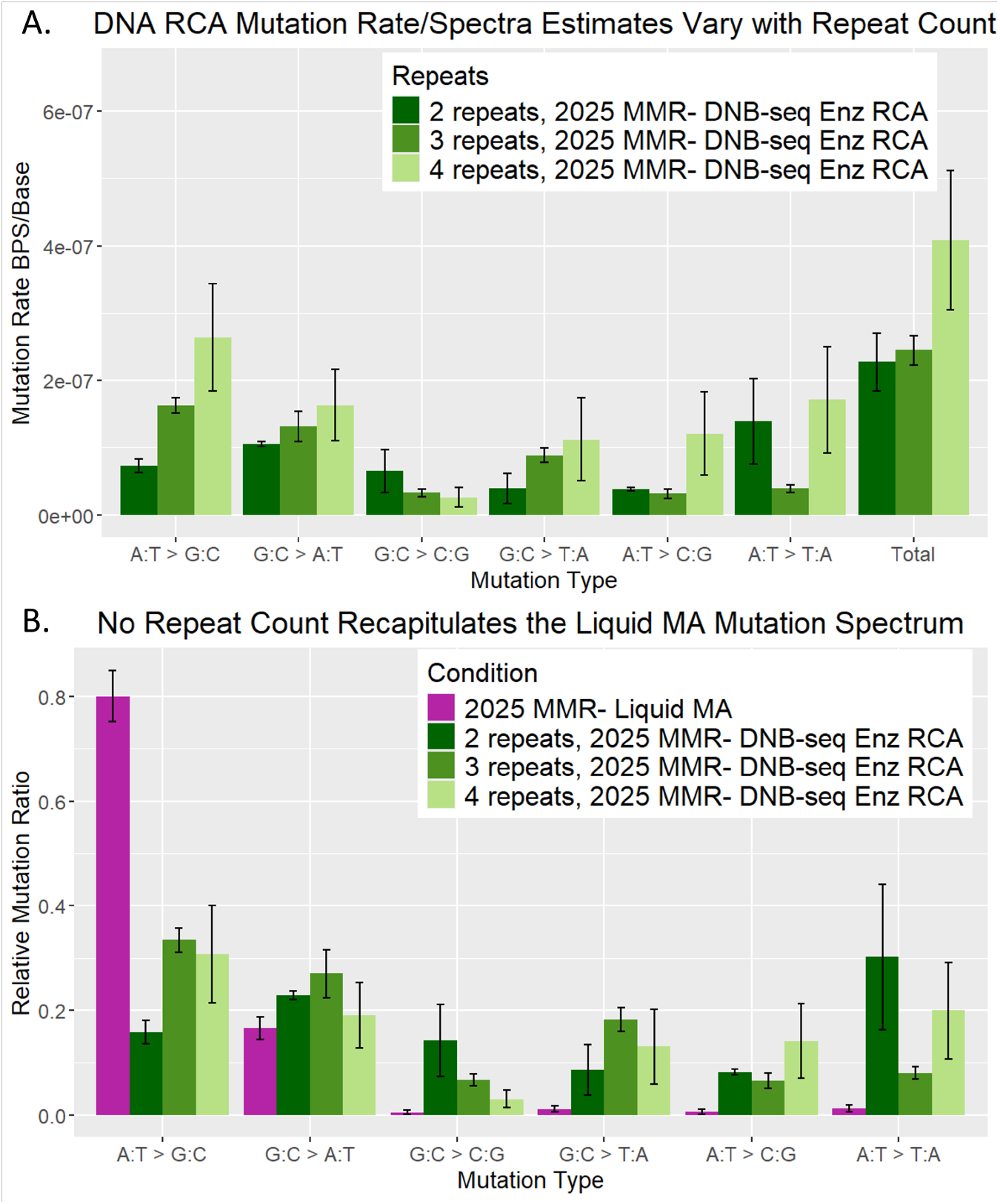
Circle-seq mutation rate and spectra vary with repeat count for MMR-*E. coli*. A. The use of 2, 3, or 4 repeats are compared for their mutation calls, which result in mutation rate and spectra estimates. B. A comparison of mutation ratios extracted from A, relative to the mutation spectrum by ratio for the 2025 MMR-Liquid MA.

As to why the 4-repeat sample calls a higher mutation rate (Figure 2a) and a less compelling mutation spectrum (Figure 2b), we surmise that a paucity of data is a potential cause. (Supplemental Table 1)

**Table 1:** Summary statistics for *N/S, dN/dS*, and genic/intergenic metrics in liquid culture w/ DNB-seq. Relative to random simulations of the observed mutation spectra, the observed locations of the mutations are sometimes significantly different from the null hypothesis. The mutation spectra of DNA Circle-seq 3 repeats and of the liquid MA were used to simulate random mutation in the *E coli* genome for 50 iterations. A mutation count roughly 540 mutations, equivalent to the number of mutations which give rise to the dnarca3 repeats mutation spectrum was generated in each iteration. *N/S* = non-synonymous/synonymous mutations; *dN/dS*; observed/expected(random sim) *N/S* ratio.

|  | LiqMA |  | Cirseq-2 |  | Cirseq-3 |  | Cirseq-4 |  |
| --- | --- | --- | --- | --- | --- | --- | --- | --- |
|  | sim | obs | sim | obs | sim | obs | sim | obs |
| N/S_mean | 1.88 | 2.13 | 2.95 | 2.51 | 2.35 | 1.39 | 3.44 | 1.0794 |
| sd | 0.21 |  | 0.31 | 0.50 | 0.23 | 0.43 | 2.41 | 0.39 |
| dN/dS |  | 1.13 |  | 0.85 |  | 0.59 |  | 0.31 |
| z |  | -6.69 |  | -7.11 |  | -8.64 |  | -0.35 |
| pval |  | 8.93E-11 |  | 4.64E-12 |  | 2.06E-17 |  | n/s |
| Genic/Inter | 7.97 | 1.14 | 7.79 | 9.86 | 7.59 | 5.26 | 7.82 | 3.69 |
| sd | 1.16 |  | 1.29 | 7.74 | 0.97 | 1.51 | 4.28 | 2.00 |
| z |  | -5.73 |  | 3.80 |  | -2.57 |  | 1.87 |
| pval |  | 3.99E-08 |  | 0.00058 |  | n/s |  | n/s |

In a further assessment of the mutations called by 2, 3, or 4 repeats, Table 1 investigates the context of the mutations called. To generate a null hypothesis of mutations upon the *E. coli* genome, we took the ratios of the mutation spectrum. With these ratios, we simulated roughly an equal number of mutations to the mutation count of the DNB-seq 3 repeats result, 540 mutations, randomly hitting the *E. coli* genome. We ran the simulation 50 times, and from those simulations estimated the average and standard deviation of several parameters: non-synonymous mutation count, synonymous mutation count, genic mutation count, intergenic (between genes) mutation count. We report the ratio of non-synonymous to synonymous mutations, a ratio of genic to intergenic mutations, and calculate a *dN/dS* ratio. The *dN/dS* ratio is a measure of selection in evolutionary biology; a ratio of 1 indicates neutral evolution, indicates ¿1 positive selection, and ¡1 indicates purifying selection. MA experiments are well characterized with regard to these values.

Table 1 demonstrates a marked impact of the mutation spectrum on the above parameters. The null hypothesis of a mutation accumulation experiment is the approximation of neutral evolution, general the absence of selection. We expect a similar approximation in this liquid culture, which so closely mimics the Liquid MA experiment. For example, the *N/S* ratio null expectation is strikingly different between the mutation spectrum simulations of Circle-seq 3 repeats and the Liquid MA, due to the relative distribution of A and T nucleotides in the *E. coli* genome. The liquid MA observed N/S ratio was slightly elevated relative to the MA simulation, though not statistically significant. This suggests the potential for weak positive selection, as evidenced by the *dN/dS* ratio of 1.13. We note that the case of Cirseq, 3 and 4 repeats does not perfectly recapitulate the liquid MA; the *dN/dS* ratio suggests the presence of fairly strong purifying selection. It is unlikely that purifying selection of this magnitude actually taking place in the liquid culture growth tube; likely, this result is an artifact of biases introduced to the system. For example, it is well known that the enzyme responsible for genomic fragmentation, MNase, has a bias, preferring to digest/fragment AT base pairs relative to GC base pairs[12]. There may also be base-pair biases to ssDNA cyclization (circularization) which may impact this parameter; ssDNA cyclization is discussed later. It appears that DNA Circle-seq, while likely able to detect mutations of high frequency from cells, may have a strong bias to how sensitive the assay is depending on the nucleotide context of the mutations.

During the course of experimentation on DNA Circle-seq, numerous conditions were examined to obtain lower mutation rates, more realistic mutation spectra, higher throughput, and greater reproducibility. Progress toward increased sequencing error resolution is detailed in Figure 3a. By iterating on the same basic idea as in the Lou 2013 result, significant resolution increases are shown, between 1500-fold and 25-fold, depending on the starting point used. We note that the change between the Hiseq-2500 and Novaseq-6000 machines, our Illumina library prep protocols changed and may have influenced these results. The change to DNB-seq was especially intended to remove the variable of library prep troubles from the assay.

**Figure 3.**
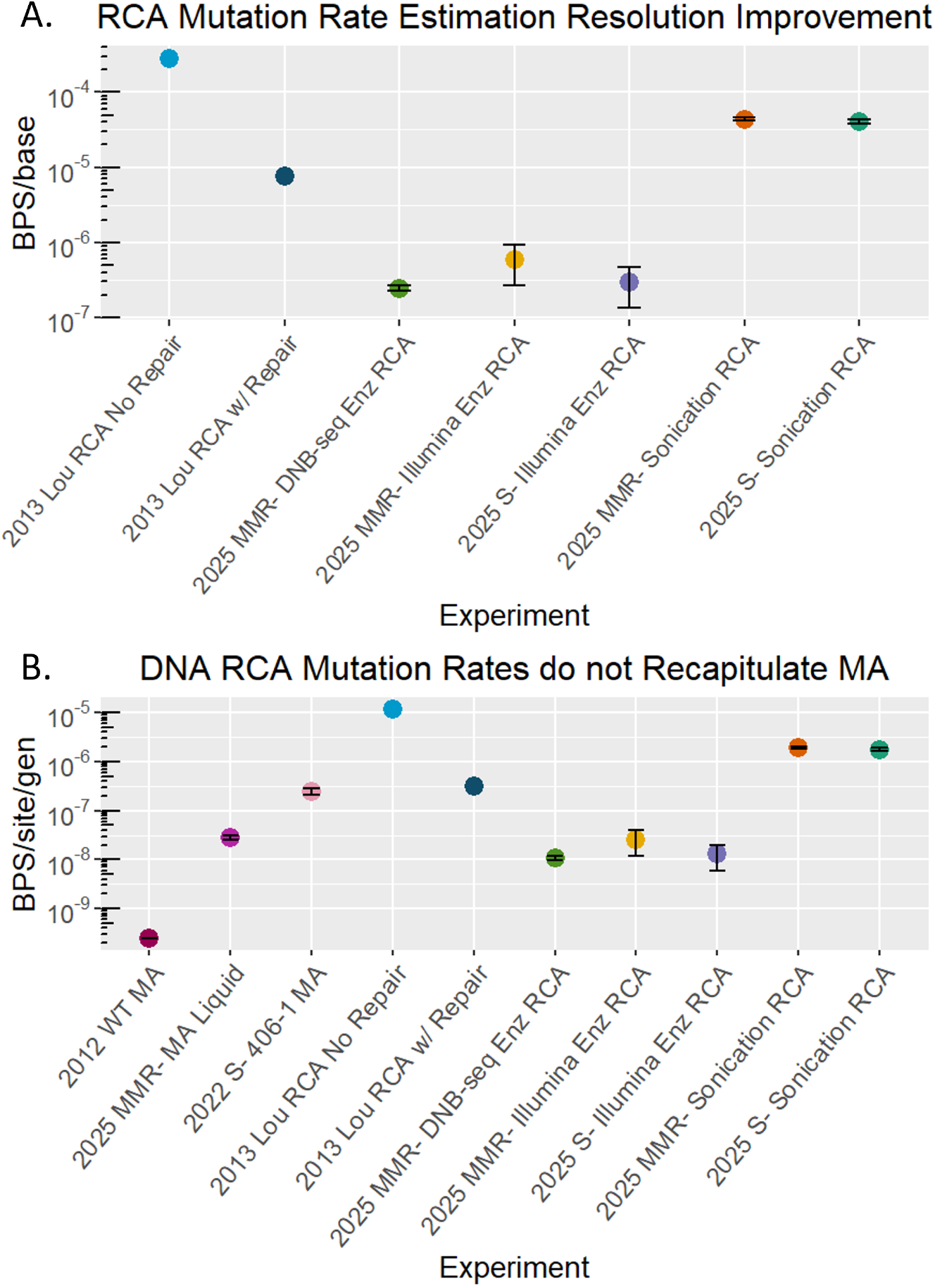
A.Circle-Seq Mutation Rate Summary. A comparison of Circle-seq mutation rate estimates of MMR-*E. coli*, in comparison to published estimates of MA experiments (left) and the original 2013 Circle-sequencing paper (Lou et al., 2013), which are based upon yeast genomic DNA. B. Conversion of the bps/base rate to bps/base/gen, and compared to bps/site/gen commonly used in MA experiments. E.c: *E. coli*. MMR-: Mismatch-Repair Deficient. Error bars: Standard Error of the Mean.

The advent of NGS-depth-based mutation queries has given rise to the measurement “mutation rate per base called” [7]. We report this number, but have made some attempt to compare mutation rate per base to the MA-experiment gold standard metric, mutation rate per site per generation (bps/site/gen)[29]. The generation count, number of cell divisions, is easily measured by CFU and log2 calculation. We note that as more bases are sequenced, more mutation calls are made, and should remain fairly constant for the first 10,000 or so mutations called. We compare the mutation rates to each other in figure 3b, where the known mutation rates of various *E. coli* strains are compared to the circle-seq estimated mutation rates on a per-cell division basis. Contrary to our expectations, we find that the mutation rate estimates are lower than the baseline MMR-liquid MA rate. It may be that we are simply missing a large pile of AT>GC mutations due to the biases of MNase or cyclization. It may also be that we are failing to remove all *in vitro* DNA damage. Regardless, we note the incongruence, and must settle by saying there are difficulties in directly comparing the MA mutation rate estimate to the Circle-seq estimate. A discrepancy between mutation rate estimating methods is not unprecedented; both the fluctuation test and MA experiment are sensitive means of detecting mutation rates; however, they cannot be exactly aligned to one another. Fluctuation test estimates converted to bps/site/gen are invariably lower than MA estimates[25, 42].

One outstanding feature of our experimentation is the impact of Covaris ultrasonication on the mutation rate estimate and mutation spectrum of MMR-*E. coli*. The paired novaseq samples arose from the exact same dna extraction test tubes, differing only in the method used for genome fragmentation. The sonicated genomic DNA produced a much higher mutation rate estimate, approaching 50-fold the samples prepared by enzymatic fragmentation (Figure 3a, Table 2). The ultrasonication used for these data points are abnormal from standard uses; to get mean fragment sizes of 80-100bp, sonication duration for the S220 Covaris ultrasonicator extended to 50 minutes per sample, from the customary 1-7 minutes suggested by the Covaris quick-guide. When a dose-response relationship is inferred, the mutational hazard of covaris ultrasonication may be on the order of 7.6 DNA damage events per 1 billion nucleotides per second. The mutation spectrum of ultrasonication is starkly different from the samples prepared by enzymatic fragmentation, showing an abundance of G:C damage in all directions; GC to AT, GC to CG, and GC to TA (Figure 4a). This is also readily demonstrated in the comparison of mutation ratios (Figure 4b). We note that all mutation rate estimates of the mutation spectrum are affected by ultrasonication; no nucleotide is safe. However, the magnitude of the deviation is far greater for the G:C base pairs (Supplemental Figure 1).

**Figure 4.**
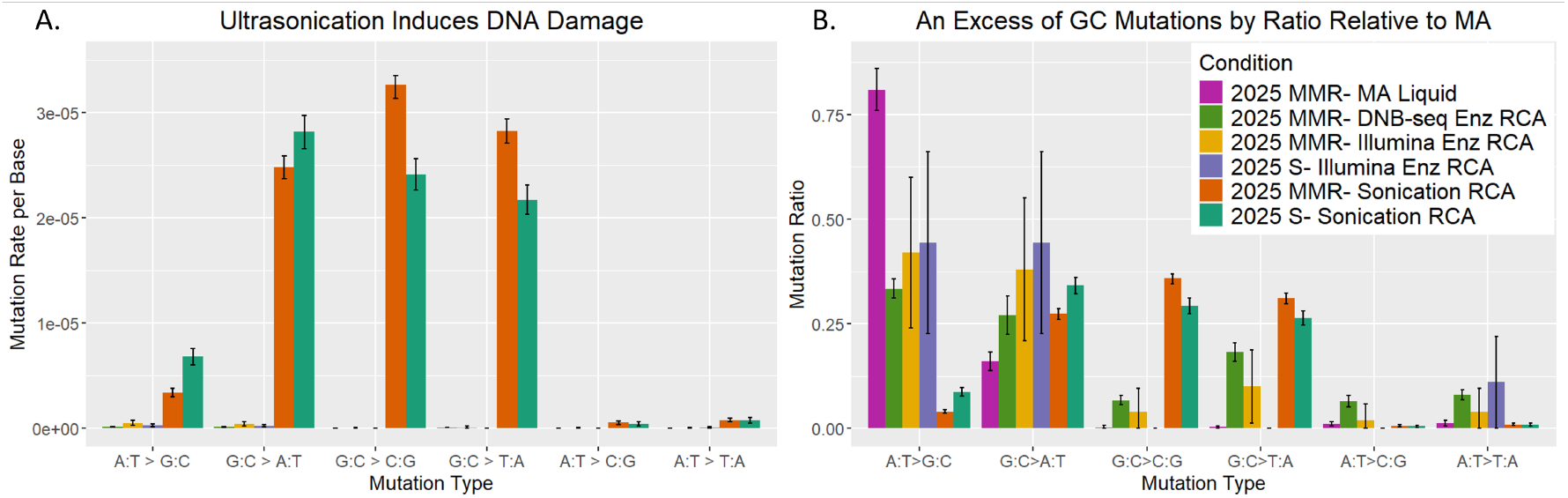
The impact of ultrasonication upon circle-seq. A. The ultrasonication mutation spectrum relative to other mutation spectra, rates. B. Ultrasonication mutation spectdrum relative to other mutation spectra, ratios. Error bars: standard error of the mean.

Repeat length has previously been mentioned as a concern and crucial variable for tweaking sequencing efficiency. Repeat length is governed largely by the distribution of fragment size generated from the genome fragmentation step. Checking repeat length is one of the first quality control measures of any experiment; and we noticed that the distribution does not quite approximate the Agilent tapestation D1000 HS fragment size distributions we generate in the library construction process. We noted a distinctive sawtooth pattern of 10bp periodicity within the fragment size distribution. The pattern became less easily discerned when the protocol changed from a 30 min DNA rolling circle amplification step to 6 h, a change that allows the protocol to use only 125ng of DNA as input. However, periodicity remains detectable (Figure 5a) among the sonicated MMR-DNA samples, and even somewhat seen in the enzymatic fragmentation samples (Figure 5b). Upon reflection, it is clear that the DNA circularization step, or cyclization step, is the source of the periodicity. Relatively more work on dsDNA cyclization has been published over the last 50 years, and in 1983 researchers noted a strong relationship between DNA twist and cyclization frequency[36] Models of dsDNA cyclization consider twist as a major factor in cyclization efficiency[21, 37, 40, 44], and it is perhaps unsurprising that RNA ligase I prefers a certain ssDNA twist as it circularizes. MNase enzymatic fragmentation of the genome appears to have more sequence bias, and results in a non-uniform distribution of fragment sizes, relative to ultrasonication. The signal has likely been depleted by the 6 hour amplification step: which fragments amplify best, and to what degree, random or biased, is not the same distribution of fragments as those which circularize. These data suggest that ssDNA twist is more important than nucleotide composition for determining fragment circularization efficiency.

**Figure 5.**
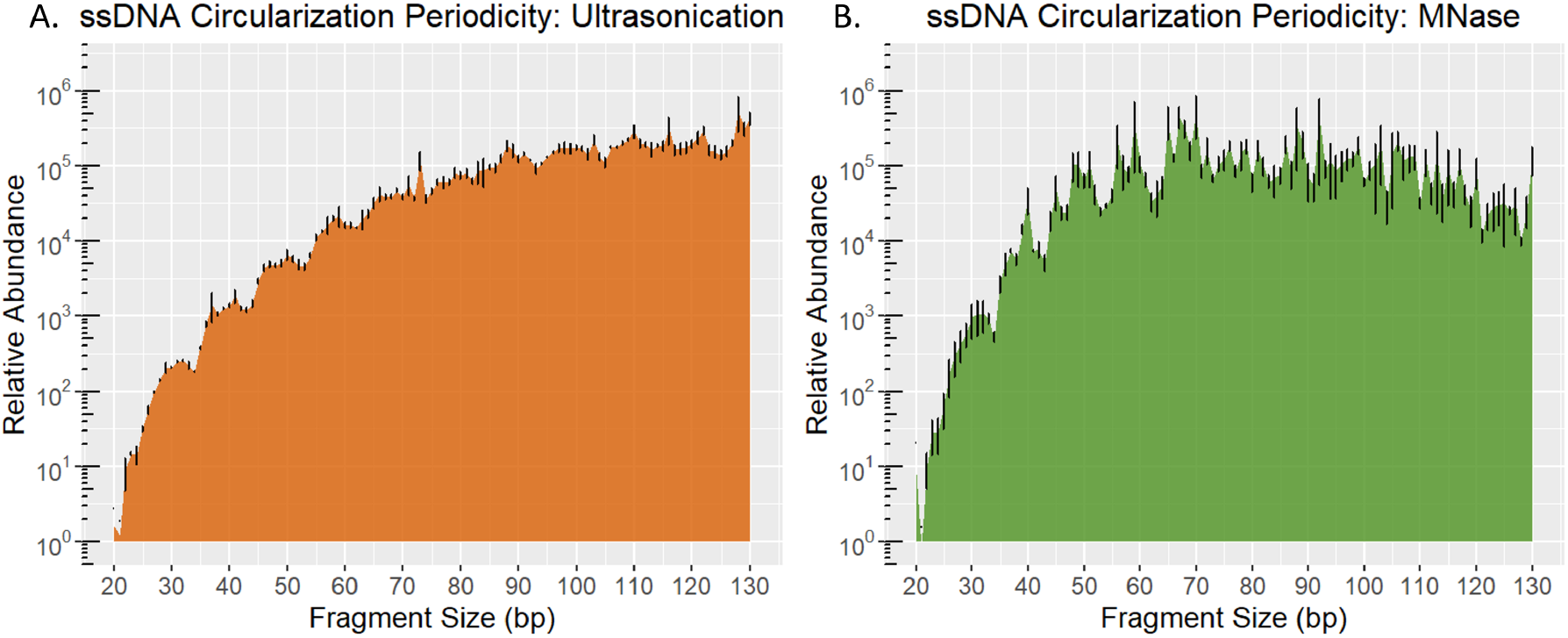
Fragment size distributions of Circle-seq. A. Relative abundance to fragment sizes of sonicated amplified MMR-*E. coli* DNA. B. Relative abundance of MNase fragmented amplified MMR-*E. coli* DNA. Error bars: Standard Error of the Mean

Amplification of the *E coli* genome, by fragment size and GC content, is biased and yet random; it is most certainly non-uniform. Figure 6 compares a general case of genome coverage across the entire *E. coli* chromosome in 1kb bins, between DNA Circle-seq and standard Illumina PE150 sequencing. The Y axis, log transformed, illustrates the magnitude of the bias, with the average sequence depth of this sample being around 100x coverage, with some regions exceeding 100,000x coverage. Some genomic regions reproducibly receive low or high coverage from DNA Circle-seq; where other regions have no such pattern.

**Figure 6.**
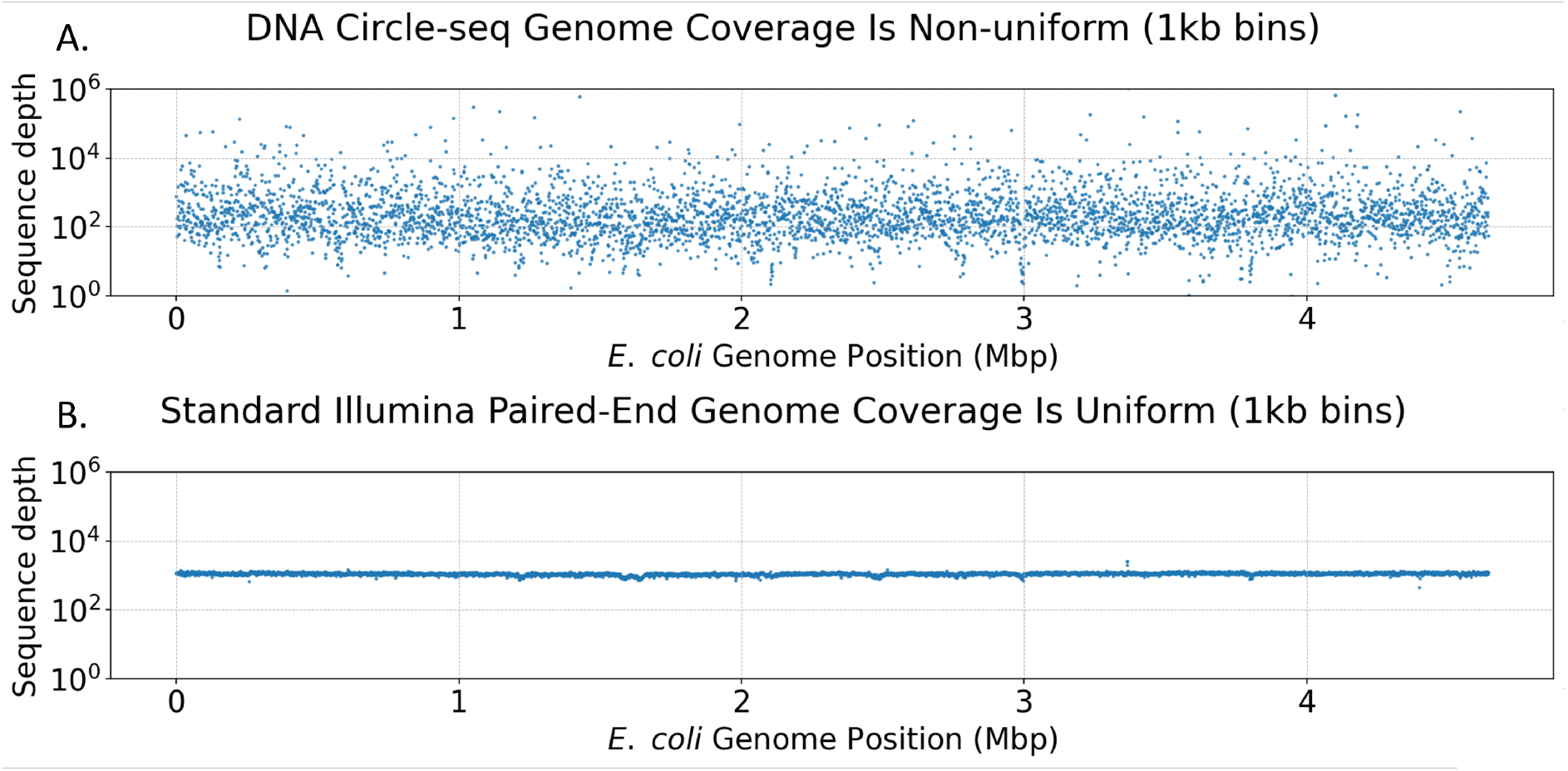
Sequence depth of reads across the *E. coli* Genome. A. Coverage depth spans 5 orders of magnitude. Each dot represents the mean sequence depth of 1000 base pairs. B. Illumina Paired-end 150bp sequencing from the Liquid MA experiment. The *E. coli* genome has been split into 4641 bins of 1000 bp.

The degree of amplification bias provides an open question about mutation calls; are the mutation calls over-represented in high coverage locations? Figure 7 addresses this concern, and suggests that the high coverage regions are not responsible for more mutation calls than other parts of the genome. Overall, with sufficient sequencing depth, or with more input DNA and relatively less amplification time, this protocol might be usable to cover similar percentages of the human genome. However, even within the *E. coli* genome there is some instance of repetitive DNA which causes issues in analysis; the problem will be far greater in the human genome. Ultimately, the short repeat length is somewhat less attractive, no matter the sequencing cost. However, the reported reduction in presumably *in vitro* DNA damage should translate to other protocols seeking to detect DNA mutations.

**Figure 7.**
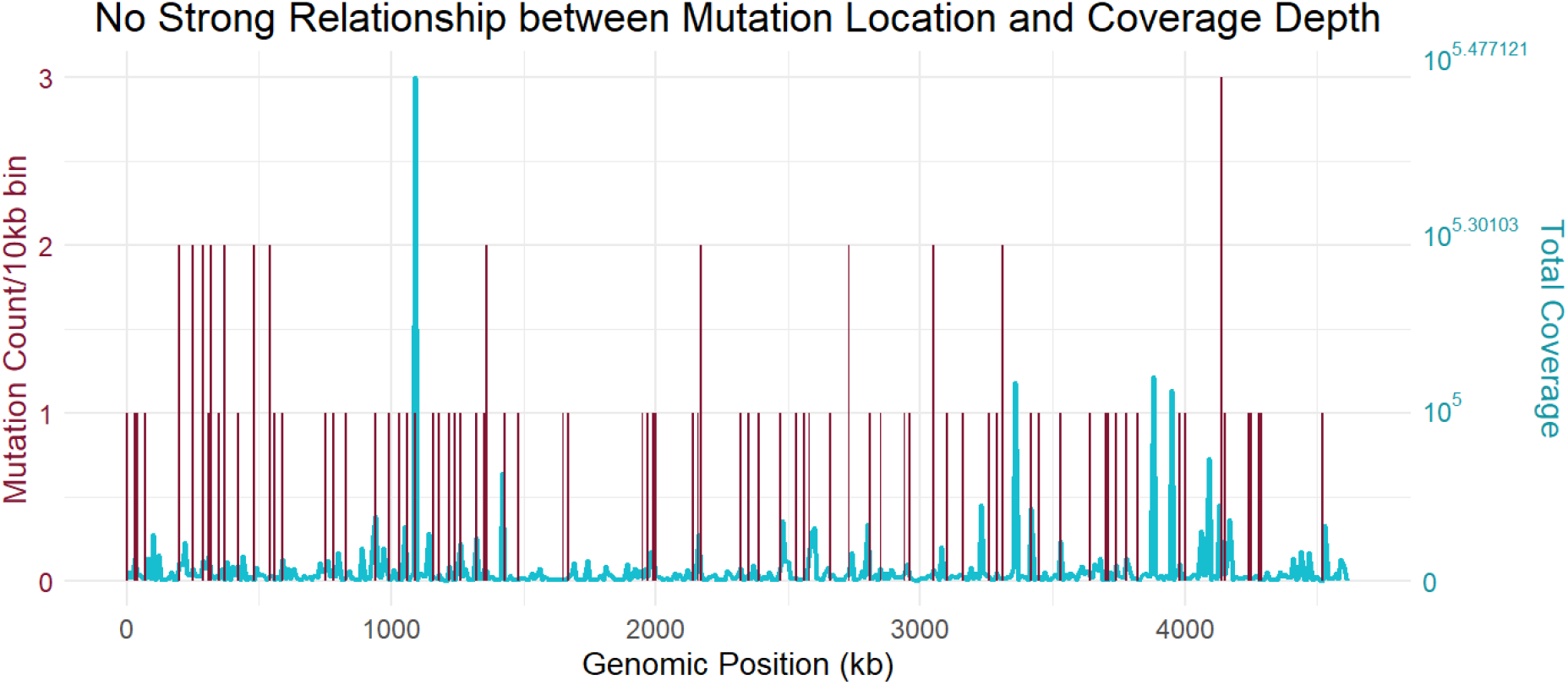
Relationship between mutation call location and sequencing depth peaks. Y axis 1: summed mutation calls from the 3 DNB-seq samples. Y axis 2: summed sequence depth from the 3 DNB-seq samples.

## 4 Discussion

We demonstrate an updated version of Circle-sequencing, which has increased the technique’s sequencing resolution by 50-fold or 1000-fold, depending on the baseline used of the original 2013 paper published by Lou et. al. Circle-sequencing is significantly improved, though as yet proves incapable of entirely resolving the mutation spectrum of MMR-*E. coli* in liquid culture. Circle-sequencing in our hands has a resolution floor of about 2 × 10^−7^ per base. The protocol has been adapted to require only 125ng of input DNA, can run on PE150 or PE250 sequencing platforms, and outputs in excess of 1 microgram of DNA from which PCR-free libraries may be constructed. We quantify the distinct mutational hazard of sonication upon DNA library preps, finding a mutational hazard on the order of 1 in 100,000 bases.

The perspective that an imbalanced mutation spectrum provides from MMR-*E. coli*, compared to a relatively balanced mutation spectrum of many WT organisms, has been incredibly helpful toward diagnosing the quality of circle-sequencing error detection. We note that this may also helpful for other sequencing techniques used to measure low mutation rates, such as laser-capture micro-dissection, duplex-sequencing, barcode sequencing, or any as-yet-untested sequencing technique that attempts to measure low mutation rates by direct detection in high-throughput sequencing datasets.

Even absent total resolution power in a technique, Circle-sequencing may prove a useful means of measuring somatic mutation burdens in human tissues and cancer. Mitochondrial heteroplasmy[19] would be relatively easy to detect. Further experiments toward this goal will be helpful in understanding this protocol’s attributes and limitations. That a single DNA extraction sample, without any laser-capture micro-dissection, or other intricate splitting scheme, requiring only 125ng, can provide information about mutation burden is potentially quite helpful to our understanding of biological mutation phenomena. For example, we expect circle-sequencing would detect rare variants within a tumor sample that are missed by conventional sequencing techniques, without significantly adding to cost. The elegance of the experimental design is likely superceded by some version of the Vijg group’s assay[30] which can also detect *in vitro* DNA damage by a convincing mechanism distinct from true mutations, while circle-seq cannot.

The circle-sequencing protocol has been made cheaper, with less reliance on expensive proprietary consumables, for example by dropping “ssDNA Ligase”, a sonication step, and eschewing “Fragmentase” as a means of fragmenting dsDNA. A practical lifetime-supply of MNase can be purchased for about $81.00 at the time of this writing. The fragmentation assay is highly reproducible and is useful over diverse DNA GC contents. The full protocol (Supplemental Information) has been reproduced across diverse sequencing platforms and with different fragment size inputs.

Circle-sequencing’s resolution floor of 2 × 10^−7^ per base is likely not the absolute resolution floor of the technique. Lou et al. demonstrated in 2013 the potential of DNA repair enzymes to reduce sequencing error rates; this technique would likely also apply to this protocol, though we did not test it. The resolution floor’s limits are readily visible: Circle-sequencing’s ability to resolve the differences between MMR-and S-lines is weak, as seen in Figure 3. MMR- and 406-1 *E. coli* strains have previously been reported to have a 10-fold difference in mutation rate, by mutation accumulation experiment. Though circle-sequencing does find a difference between the two strains in the correct direction, it is not quite a difference of 2-fold. For this, we can only speculate that the *in vitro* DNA damage signal, or other biases in the system are confounding the protocol’s ability to fully resolve this difference. Regardless, we are able to report reproducibility of the experiment across time and sequencing platforms. With sufficient sequencing depth, the error bars from sample to sample have been limited; for example, in the DNB-seq sequencing runs.

We note that there is likely a lower bound to how short a fragment can be, at which point rolling-circle amplification fails. If the mean fragment size drops below 50 bases, we have noticed somewhat reduced yield from the experiment given an equal amount of DNA input. Further, the percentage of reads with repeats drops from 50-80% to as low as 9%; this is likely due to the existence of some fragments larger than 200 base pairs that circularize. Circularized DNA larger than 200bp, be they single fragments or oligomers, will escape the exonuclease digests, and will amplify with far greater efficiency than fragments of 30bp. Even if no large fragments can be detected on a tapestation, picogram amounts can offset the efficiency of an experiment. Running a gel is an acceptable means of avoiding this problem, although it is also possible to simply aim for an ideal repeat length of about 60-80bp. A Bluepippin machine may prove useful in refining the distribution.

That ultrasonication is a 100-fold mutagen is unexpected. The original intent of using sonication was to find a less-biased means of fragmenting genomic DNA. There is some circumstantial commentary throughout the DNA sequencing field that ultrasonication is somewhat mutagenic[11**?**], but we have not found any reports that so clearly demonstrate the magnitude and specific mutation spectrum. In consideration of COSMIC mutation signatures[14], it is possible one of the mutation signatures detected by the research has a component of ultrasonication, or should be added. We hasten to add that ultrasonication/cavitation is used in the health industry as a form of liposuction, as a means of breaking up adipose cells. However, the frequency of the Covaris ultrasonicator and human applications appear to be entirely different from one another. To what degree any cavitation may cause DNA damage and not cell lethality is unknown but appears to be a low risk; and over decades of use, no link between ultrasonication cavitation and cancer has been reported. The authors also imagine that massive fragmentation of the genome is a far greater concern to cells than a relatively modest increase in DNA damage. We anticipate the usefulness of information about the DNA damage potential of ultrasonication is primarily toward considerations of which library prep to use when searching for somatic mutations.

## 5 Acknowledgements

Author S.B. wrote the manuscript, lead experiments, participated in analysis, and experimental design. J.F.G. reviewed the manuscript many times, lead analysis, participated in experimental design. L.R. and H.R. assisted in experiment execution and troubleshooting for years. M.L. participated in experimental design, manuscript review, and obtained funding for the project.

We would like to thank Dr. Megan Behringer, Dr. Wei-Chin Ho, and Sam Miller for their recurrent feedback over the years of Lynch lab meetings. Figure 1 was created with the assistance of Biorender.com.

